# Adult Drosophila Muscle Morphometry through MicroCT reveals dynamics during aging

**DOI:** 10.1101/587733

**Authors:** Dhananjay Chaturvedi, Sunil Prabhakar, Aman Aggarwal, K VijayRaghavan

## Abstract

Indirect Flight Muscles (IFMs) in adult Drosophila have served as a valuable model for studying muscle development. In terms of function, they provide the key power stroke in adult insect flight. Variability in their architecture including of fiber numbers, shape and arrangement may provide insightful clues into adult muscle function. Conventional histological preparations in imaging techniques severely limit exact morphometric analysis of flight muscles, thereby impeding causal or correlative studies between muscle morphology and function. In this study we employ MicroCT scanning on a tissue preparation that retains muscle morphology under homeostatic conditions. We use this method to deliver precise measurements of a subset of IFMs, the Direct Longitudinal Muscles’ (DLMs) size and shape, in male and female Drosophila and changes therein, with age. Our findings reveal several unexpected characteristics of muscle fibers. We also demonstrate application to other insect species making it a valuable tool for histological analysis of insect biodiversity.

**Significance Statement:** Adult Drosophila muscles serve as models of homeostatic muscles. Accurate analysis of their form and function is key to understanding affects of genetic and physiological states on them. Recording adult muscle shape and volume has so far depended on protocols that inevitably distort tissue. Here, we use a MicroCT scanning based method that delivers changes in shape, size and organization between males and females, with time. This method is a significant step forward in recording muscle structure *in situ* with applications across species.

## Introduction

Dorsal Longitudinal Muscles (DLMs) are a subset of adult insect flight muscles(1). DLMs are the only reported large Drosophila muscle group, which shares myofibrillar architecture with mammalian skeletal muscles. Despite significant differences in modes and frequency of activation from mammalian counterparts, they have been exceedingly informative in the understanding in vivo principles of muscle development such sarcomere formation(3).

DLMs are arranged in sets of six muscle fibers, one beneath the other, running anterior to posterior on either side of the sagittal midline, in the dorsal thorax of adult Drosophila (Fig 1a.). These large syncitia arise during pupariation by the fusion of Adult Muscle Progenitors(4) to remnants of larval body wall muscles that escape histolysis, called templates. 8hrs after puparium formation, histolysis of larval body-wall muscles surrounding three templates on either side of the midline is complete. Two syncitia result from the splitting of each template by 20hrs apf, which is how six fibers are seen on either side of the midline in adults(5). Zero or more than one splitting events lead to aberrant, astereotypical DLM numbers. Until adult flies eclose from their pupae, muscles undergo phases of rapid growth in larvae, histolysis in pupariation, followed by rapid growth again. Post pupariation, muscles attain stable structures without catastrophic events like histolysis. This state is ideal to model homeostatic muscle tissue.

**Fig 1.**
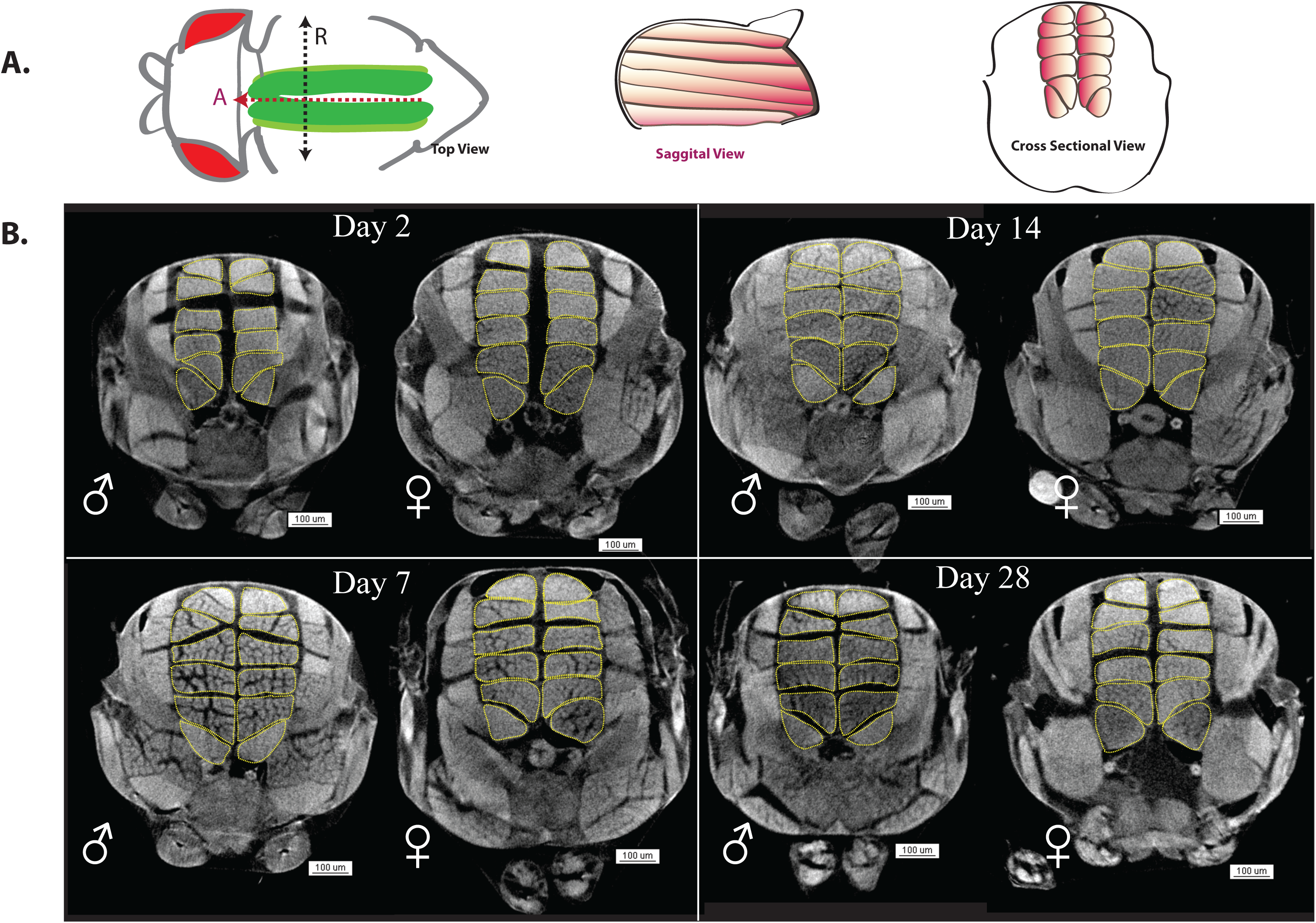
Gross changes in male and female Adult Drosophila Longitudinal Muscles over time. **(A)** Schematic of Dorsal Longitudinal Muscle (DLM) positioning in the adult thorax. In Top view, DLMs (Green) run along the anterior -posterior axis (red dotted line, arrowhead indicates anterior ‘A’) in the thorax. Here the cuticle is imagined to be transparent to highlight their position relative to the head and within the thorax. The black dotted line describes the left right axis, running between the wing hinge. ‘R’ denotes the animal’s right hand side. Orthogonal to the plane of top view, through the red dotted line (sagittal view), six muscle fibres (orange) run anterior to posterior arranged in a dorsal to ventral fashion, in one hemithorax. Orthogonal to the plane of top view, through the black dotted line (cross sectional view), six DLMs are arranged in the thorax, on either side of the midline as shown. **(B)** Representative cross sections of whole thorax MicroCT scans of male(♂) and female (♀) flies at days 2, 7,14 and 28 post eclosion (p.e.). DLMs are outlined in each cross section with yellow dotted lines. Scale bars: 100µm; n=14-21 per sex per time point.

Muscle development in Drosophila has been studied for decades. These studies have revealed *in vivo* molecular insights into muscle development. Investigations into the interaction between neurons and muscles at NMJs have proved valuable. Much of these studies have been carried out in larvae or pupae. Ease of staining muscles with dyes, fluorescent or otherwise due to their relatively small dimensions at these stages, makes them very amenable to microscopy. Specifically, examining protein localization within cells, cellular organization and structure, and positioning of cells in the context of neighboring tissue types has been achieved with great precision. However, examination of these parameters in adult Drosophila muscles with increased Z axis coverage is limited by technical challenges such as antibody penetration and tissue clarity. Often, circumventing these issues through the use of clarifying agents or detergents leads to altered muscle morphology. For instance, the use of alcohols for fixation and as clearing agents leads to severe tissue shrinkage, thereby not reflecting tissue structure faithfully.

Cytological details such as sarcomeric structures, nuclear arrangement, muscle-tendon and nerve interactions in Adult Drosophila muscles have been examined with great resolution through confocal microscopy. Cuticle opacity has remained a significant obstacle in detecting signal from structures and cell populations beneath the surface of the animal. Further, even sufficiently strong fluorophores fail to provide three-dimensional information on samples exceeding ∼30 microns along the Z axis. Therefore, current confocal microscopy protocols have found limited applications in three-dimensional modeling of whole muscle tissues *in situ*.

Current understanding of DLM morphology relies on immunohistochemistry from dissected tissue. Dissection is by definition invasive and inevitably changes the morphology of tissue to some degree. Accurate DLM morphological analysis *in situ* is made impossible in genetic conditions where DLMs are already malformed and fragile. These genetic conditions most likely are the most accurate Drosophila models of human adult onset muscle pathology. Therefore, accurate, quantitative measurement of adult Drosophila musculature is much required. Non-invasive techniques to visualize internal structures at high resolution, while being practical have been recently explored.

MicroCT analysis has proven to be a sturdy bridge between high-resolution imaging and depth of visualization. Three-dimensional visualization of tissue *in situ* is proving critical in studying tissue repair and degeneration. To this end, various fixation methods and contrasting agents have been applied to different tissues and species. Recent studies have explored several contrasting agents to characterize the anatomy of several insect species. Crucially, several of these sample preparation methods involve ethanol and high concentration of salts for fixation and storage of samples(6-8). A mere visual comparison of DLM morphologies between these preps and Immunohistochemical preps reveals compromised morphology, probably due to employment of dehydrating agents, salts and fixatives that cause shrinkage. A MicroCT scanning protocol where internal soft tissue morphology is corroborated by IHC dimensions, is much needed.

To address this need, we adapted staining with Lugol’s solution in phosphate buffer saline in order for DLMs to retain morphology as observed with conventional Immunohistochemistry protocols for DLMs. Modifications to sample immobilization during scanning have quickened the process and clarified resulting images.

With this method, DLM morphology in cohorts of male and female fruit flies up to four weeks post eclosion was measured with 8um^3^ resolution.

We have found not only consistent increases in volume, but also changes myofiber fascicle arrangement with individual DLMs. These newly discovered changes in form motivate further enquiry into changes in muscle function as well as intramuscular organelle distribution with aging. Further, we can now accurately measure deviations from wildtype *in situ* DLM morphology in case of muscle dystrophy models in Drosophila.

## Results

### MicroCT scanning of Drosophila thoraces reveals muscle structure *in situ*

We employed MicroCT scanning to visualize thoracic muscles *in situ*. All fixation and staining steps maintained osmotic conditions as close to physiological as possible. Our contrasting agent of choice was modified Lugol’s solution (1% I_2_, 2% KI in PBS). Iodine uptake by muscles has previously demonstrated (9). All scans were collected on Bruker’s Skyscan 1270 machine at 0.5μ resolution.

Video 1 shows a 3D MicroCT scan thorax of a two day old adult. Surface structures such as the chitin exoskeleton, halteres, macrochaetae and microchaetae can be seen very clearly. Muscles within the thorax including Direct Flight Muscles (DFMs), Dorso-Ventral Muscles (DVMs) and DLMs clearly show a strong signal throughout their volume.

Fig.1a. describes the position of our muscle group of interest, the DLMs, within an adult thorax. We wondered if the volumes and packing of DLMs vary with age and sex. Fig.1b. shows representative thorax cross sections from MicroCT scans of males and females of 2 days, 7 days, 14 days and 28 days post eclosion.

Animals in this cohort were collected within 1-hour post eclosion. Subgroups of 10 to 20 males and the same number of females were sacrificed at each time point and prepped for MicroCT scanning.

Sexual dimorphism (larger female thoraces), in sizes is obvious from these cross sections. Also, the tendency of DLMs to pack more tightly inside the thorax over time in both males and females is apparent.

However, the separation of fascicular structures within muscle fibers was intriguing. In Video 2, separation between actin bundles running anterior to posterior can be seen in the sagittal view. As the plane of view turns coronal and moves anterior to posterior, the dark spaces between patches of signal, run continuously without abrupt shifts between coronal planes, suggesting continuous fascicles. Also, though fascicles separate considerably in the antero-posterior midline of the muscle, they meet at the anterior and posterior ends. This suggests that regions with no signal are non-myofibril filled space between fascicles. We asked if these spaces maybe filled with muscle organelles such as mitochondria and endoplasmic reticulum.

To address this possibility, DLM mitochondria and endoplasmic reticulum, along with muscle membranes were visualized in cross sections. To this end, the muscle driver *mef2*-Gal4 was used to drive mito-GFP and KDEL-RFP in backgrounds of mCD8 tagged fluorescent proteins. In Fig 2A., membrane markers mCD8GFP and mCD8 RFP can be seen around individual muscles. mCD8-GFP appears to mark membranes more uniformly than mCD8-RFP. Given that they are both driven by the same Gal4 driver and being examined in the same tissue, variation in their distribution is likely to be due to the difference in aggregation properties of the two fluorescent proteins. Myofibril fasciculation is apparent through these membrane markers, within each muscle fiber. Each fascicle appears bounded by these mCD8 tagged fluorescent proteins. Due to the invasive nature of the dissection procedure, measuring the separation of fascicles post dissection, is unlikely to reveal *in situ* arrangement, as revealed by MicroCT scans. From these immunostainings, the distribution of mitochondria between individual fibrils (mito-GFP signal); the distribution of endoplasmic reticulum (KDEL-RFP signal) and within the fascicle is clear. Taken together, data from immunohistochemistry and MicroCT scans suggest, myofibrils appear to be arranged in membrane bound fascicles within individual DLMs. The contents of the space between fascicles are not known.

**Fig 2.**
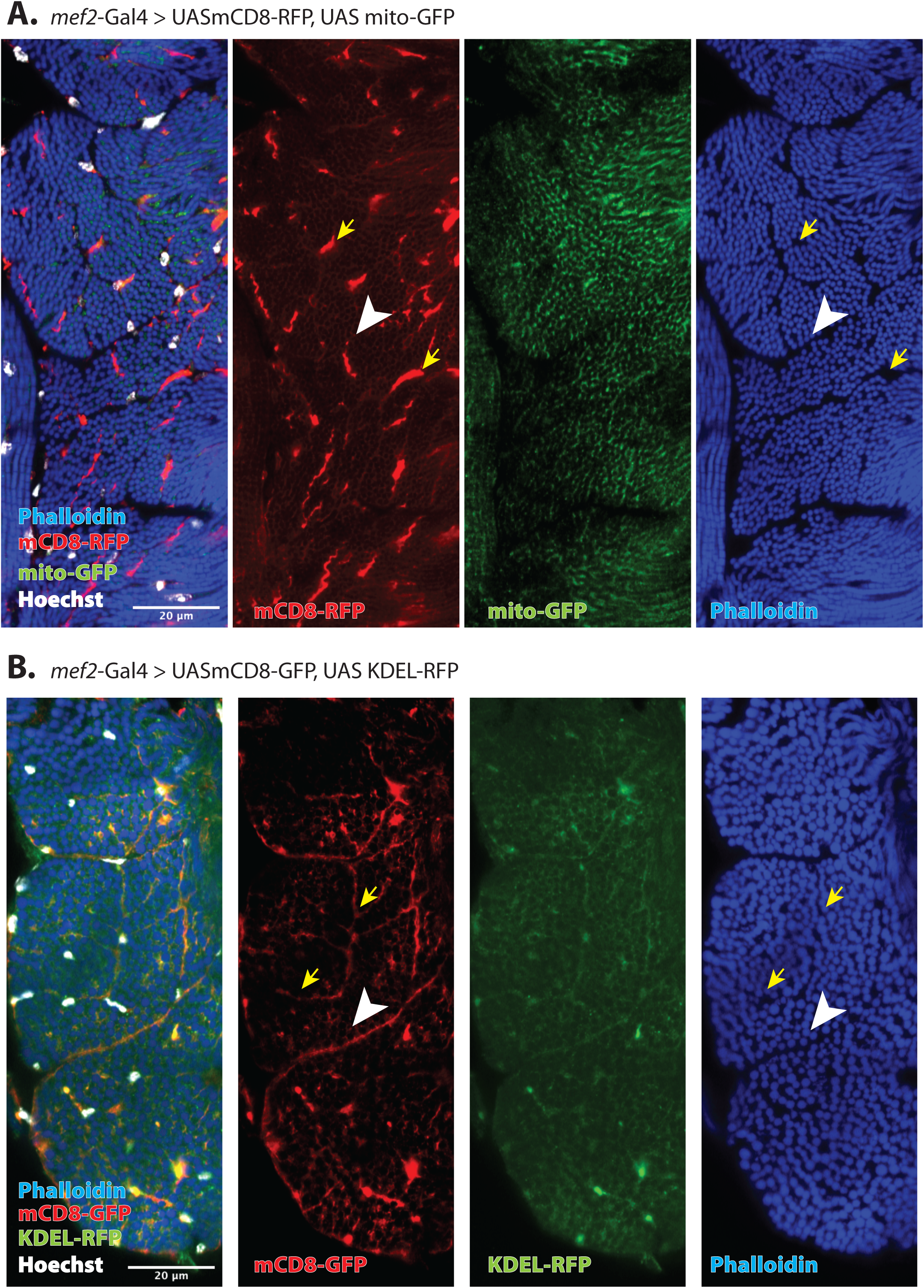
Membrane, Mitochondria and Endoplasmic reticulum distribution within DLM Fibres. **(A)** Representative cross section of DLMs from animals with muscle driver *mef2*-Gal4 driving membrane bound RFP (mCD8-RFP) and mitochondrion localized GFP (mito-GFP). Phalloidin (blue) marks myofibrils, mcd8RFP (red) marks cell membranes, mito-GFP (green) localizes to mitochondria, Hoechst (white) marks nuclei. The white arrowhead marks a boundary between two separate DLM fibres. Yellow arrowheads indicate RFP marked membrane separating two different DLM fibres and myofibril fascicles within a DLM fibre. n=7 **(B)** Representative cross section of DLMs from animals with muscle driver *mef2*-Gal4 driving membrane bound GFP (mCD8-GFP) and Endoplasmic reticulum localized RFP (KDEL-RFP). Phalloidin (blue) marks myofibrils, mcd8GFP (red) marks cell membranes, KDEL-RFP (green) localizes to endoplasmic reticulum, Hoechst (white) marks nuclei. The white arrowhead marks a boundary between two separate DLM fibres. Yellow arrowheads indicate GFP marked membrane separating myofibril fascicles within a DLM fibre. Scale bars= 20 µm; n=7

We also find that stereotype of six DLMs arranged on either side of the midline in adult thorax does not hold one out of forty animals in our wildtype *CS* culture. Six DLM fibers in each hemithorax arise from the splitting of three muscle syncytial templates during pupariation. In Fig.S1. we show three instances of adult hemithoraces where templates failed to split during pupariation or split more than once. This denotes the background rate of this specific defect in muscle development in our wildtype stock (1:40).

**Fig S1.**
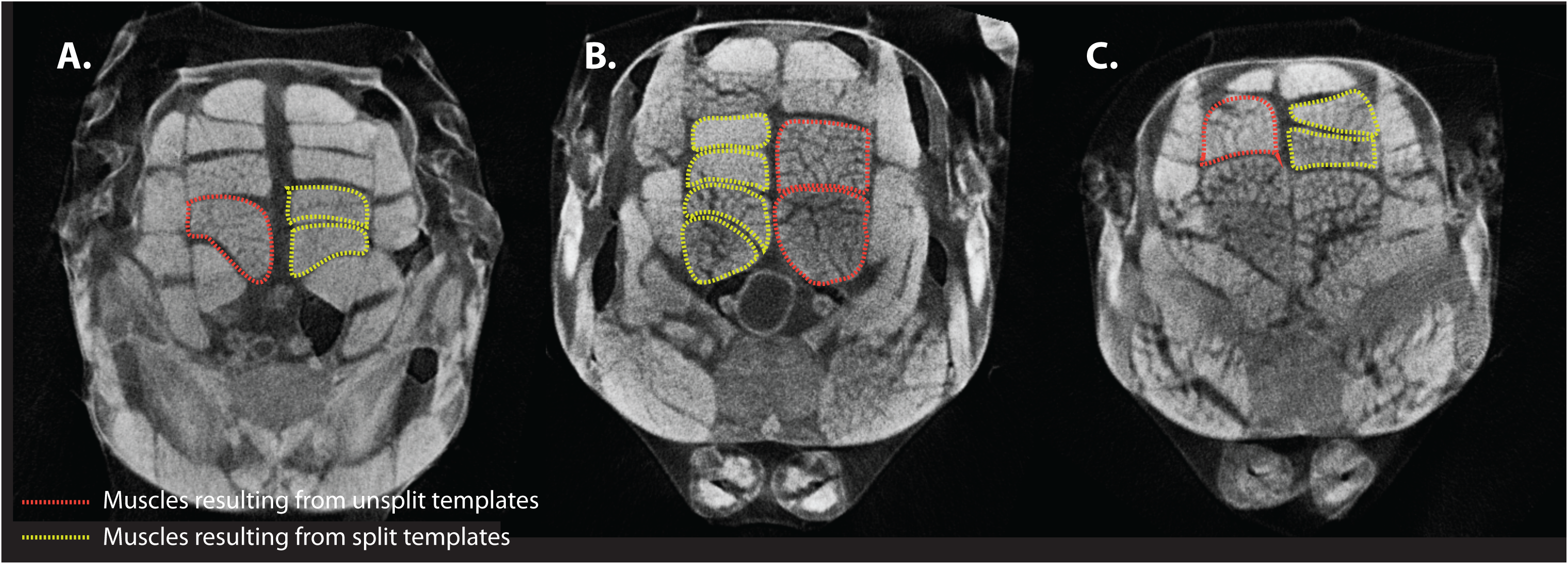
Examples of DLM fibres resulting from template splitting failure during pupariation. Cross sections of three different whole thorax MicroCT scans (A.,B.,C.). Yellow dotted lines outline DLM fibres resulting from stereotypical template splitting during pupariation. They serve as internal controls for DLM fibres, on the opposite side of the midline, outlined in red dotted lines where template splitting failed during pupariation.

### Mapping Individual Muscle Fiber Volumes over time

Thoracic musculature can be cleanly visualized from our MicroCT scans. We focused DLM fibers for our morphometry using Bruker’s CTan software volumes of individual muscle fibers. Briefly, regions of interest (RoIs) outlining the boundaries of individual muscle fibers are marked manually in three planes along the length of the muscle. The volume of each fiber thus gets interpolated. This measurement specifically gives the volume of a muscle fiber inclusive of regions that lack signal.

In Fig.3A., DLM nomenclature is described in cross-section. The dorsal most DLM is called “a” and ventral most muscle is called “f”. Which hemithorax of the animal the fiber belongs to referred to by superscripting R (right) or L (left). Table 1 lists average volumes measured at 8μ^3^ resolution for each DLM for males and females at different adult ages with observed standard deviations.

**Fig 3.**
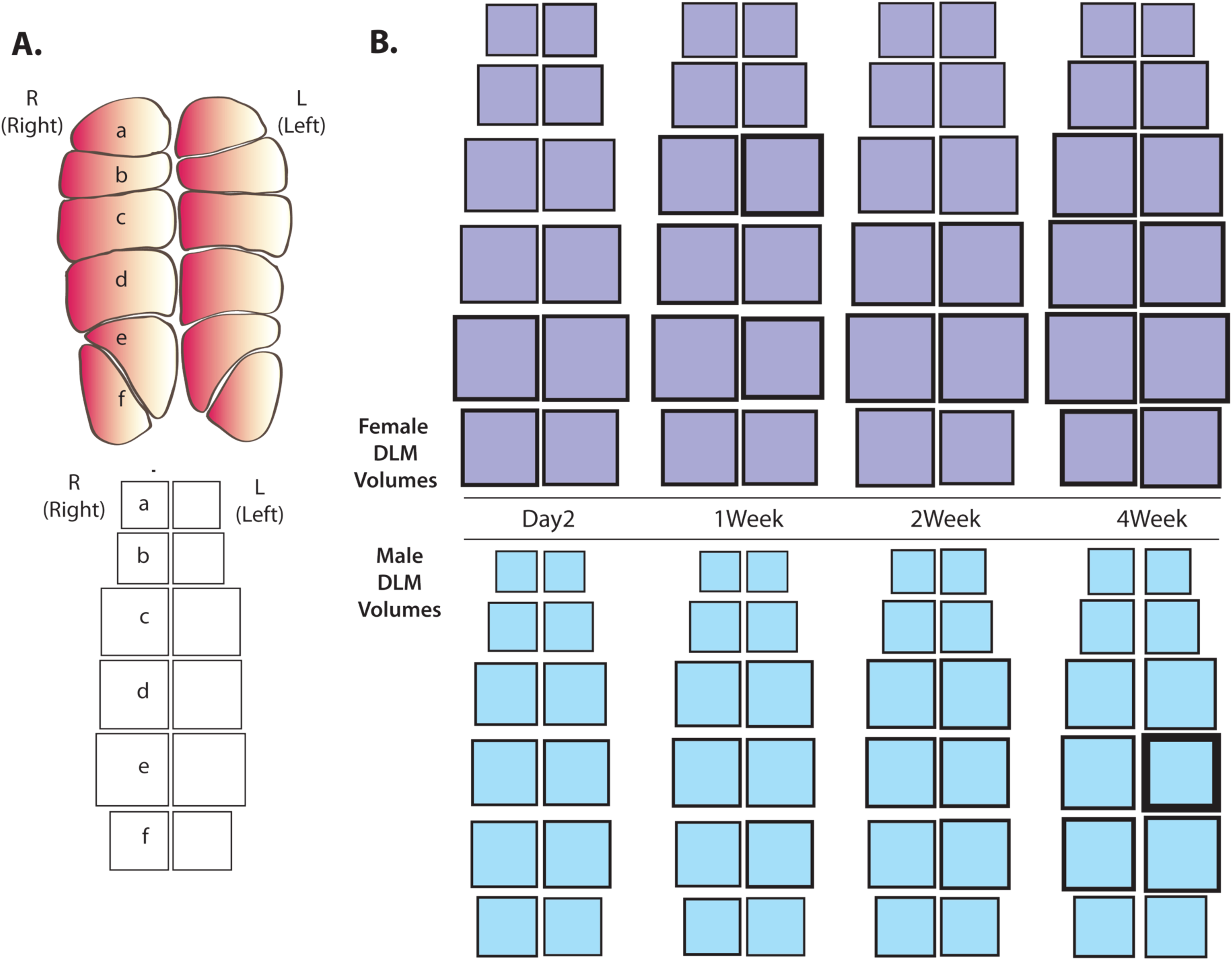
DLM Volume mapping for males and females post eclosion. **(A)** Schematic of DLM nomenclature. DLM fibres located in the animal’s left hand hemithorax are marked L and those on the animal’s right hand hemithorax are marked L. Individual fibres are marked ‘a’ to ‘f’ arranged in a dorsal to ventral manner. The lower panel describes a volume map of one full set of DLMs from one animal. The volume of each muscle is represented by a square of proportional area. Each square is arranged in the map, corresponding to the position of the represented muscle, in the thorax. **(B)** Average individual muscle fibre volumes mapped as described in (A), for females (purple) and males (blue) at days 2, 7,14 and 28 post eclosion (p.e.). The thickness of the border of each square represents half the standard error of mean in volume for that muscle, in that group, at that time point. n=14-21 animals per sex per time point.

**Table 1.**

| Table 1. | Average Volume (μ³) |  |  |  | Standard Deviation (μ³) |  |  |  |
| --- | --- | --- | --- | --- | --- | --- | --- | --- |
| Females | Day 2 | Day 7 | Day 14 | Day 28 | Day 2 | Day 7 | Day 14 | Day 28 |
| a <sup>L</sup> | 4660242.86 | 4865094.12 | 5159777.78 | 4668171.43 | 935214.35 | 729389.38 | 676056.93 | 544729.10 |
| a <sup>R</sup> | 4428057.14 | 5002994.12 | 5054944.44 | 4934014.29 | 581075.43 | 617658.01 | 551736.01 | 655400.51 |
| b <sup>L</sup> | 5912700.00 | 6781582.35 | 7055888.89 | 7438000.00 | 1002600.10 | 997585.47 | 737796.87 | 1167167.77 |
| b <sup>R</sup> | 6042550.00 | 7031305.88 | 6926727.78 | 7497400.00 | 717299.45 | 1132261.68 | 782864.47 | 1168091.47 |
| c <sup>L</sup> | 8610100.00 | 10505729.41 | 10145372.22 | 10903264.29 | 766682.94 | 3790823.40 | 1091783.74 | 1727204.83 |
| c <sup>R</sup> | 8921164.29 | 10124611.76 | 9502522.22 | 11247471.43 | 830203.66 | 1887062.79 | 1226718.26 | 2033314.02 |
| d <sup>L</sup> | 10027985.71 | 10187929.41 | 11421200.00 | 11444067.86 | 1240880.81 | 1521182.12 | 2060163.53 | 2375259.91 |
| d <sup>R</sup> | 10033935.71 | 10467247.06 | 11110611.11 | 12413707.14 | 966102.93 | 2153768.38 | 2020978.74 | 2004953.10 |
| e <sup>L</sup> | 12211642.86 | 10701758.82 | 12251255.56 | 12767842.86 | 1628844.33 | 2935291.71 | 2653294.70 | 2653294.70 |
| e <sup>R</sup> | 11769357.14 | 12011570.59 | 12746944.44 | 13403628.57 | 1043189.74 | 2279769.57 | 2227002.43 | 2227002.43 |
| f <sup>L</sup> | 9417900.00 | 9200400.00 | 9063627.78 | 10468700.00 | 1298537.66 | 1508967.70 | 1288052.33 | 1753665.48 |
| f <sup>R</sup> | 9661257.14 | 9433458.82 | 10087033.33 | 9177000.00 | 1798956.87 | 1798956.87 | 1462735.55 | 2588732.78 |
| Avg. Total DLM Volume | 101696892.86 | 106313682.35 | 110525905.56 | 115854158.82 | 5709912.22 | 9653834.64 | 11478726.51 | 28523041.01 |
|  | Average Volume (μ³) |  |  |  | Standard Deviation (μ³) |  |  |  |
| Males | Day 2 | Day 7 | Day 14 | Day 28 | Day 2 | Day 7 | Day 14 | Day 28 |
| a <sup>L</sup> | 2803093.33 | 2827637.50 | 3387014.28 | 3446918.75 | 383212.93 | 349094.30 | 765913.13 | 660614.75 |
| a <sup>R</sup> | 2866126.66 | 2752306.25 | 3207780.95 | 3541512.50 | 364171.13 | 400067.14 | 621885.19 | 514544.55 |
| b <sup>L</sup> | 4094906.66 | 4305400.00 | 4431438.09 | 4842850.00 | 488253.37 | 529423.66 | 809520.40 | 751109.61 |
| b <sup>R</sup> | 4028926.66 | 4354087.50 | 4613238.09 | 4843056.25 | 488253.37 | 524686.18 | 827511.65 | 769067.72 |
| c <sup>L</sup> | 6527500.00 | 7346200.00 | 8047928.57 | 8395881.25 | 701169.51 | 1023204.90 | 1629740.93 | 1105113.85 |
| c <sup>R</sup> | 6504166.66 | 7117368.75 | 7759990.47 | 7846768.75 | 770867.55 | 906305.07 | 1369926.16 | 1180070.62 |
| d <sup>L</sup> | 7167486.66 | 7749181.25 | 7789614.28 | 8714950.00 | 936796.96 | 1032275.20 | 1793874.29 | 8714950.00 |
| d <sup>R</sup> | 7203406.66 | 7743687.50 | 7994771.42 | 8793037.50 | 831662.05 | 1175830.90 | 1706509.61 | 1696541.42 |
| e <sup>L</sup> | 7513286.66 | 7384900.00 | 7977447.61 | 9275093.75 | 950729.03 | 1394573.60 | 1702713.38 | 2033749.84 |
| e <sup>R</sup> | 7341973.33 | 7075225.00 | 7504938.09 | 8466231.25 | 782353.23 | 803162.08 | 1180949.84 | 2351928.96 |
| f <sup>L</sup> | 5634726.66 | 5658843.75 | 5669319.04 | 6321412.50 | 525705.24 | 568036.19 | 984490.35 | 759107.27 |
| f <sup>R</sup> | 6058900.00 | 5359343.75 | 5980742.85 | 6114881.25 | 748863.08 | 835734.67 | 1033071.12 | 924268.34 |
| Avg. Total DLM Volume | 67744500 | 74319126.67 | 74364223.81 | 80602593.75 | 5728508.00 | 16791698.00 | 9613560.46 | 7391786.06 |

|  | Number of Measurements |  |
| --- | --- | --- |
|  | Females | Males |
| Day2 | 14 | 15 |
| Day7 | 17 | 16 |
| Day14 | 18 | 21 |
| Day28 | 14 | 16 |

Individual DLM fiber volumes of females and males over time

For an uncluttered, visual representation of individual muscle placement and normalized volumes within a thorax, we devised a volume mapping. The volume of each muscle fiber is represented by a square of proportional area rather than a one-dimensional bar on an axis. Each square representing a single DLM fiber’s volume is arranged according to its positioning in the thorax (Fig.3A.). The standard error of mean in measurements per muscle fiber from a group, is depicted through proportionate border thickness in pixels.

Fig.3B. shows volume maps of all twelve DLMs in males and females at different times post eclosion. The area of all squares is normalized to the smallest measurement in the entire dataset and then scaled to fit the canvas. The maps accurately depict the consistent difference in muscle volume of females versus males over time. The difference in relative volumes of female DLMs and corresponding male DLMs over time can be gleaned.

Notably, average individual muscle fiber volumes differ from the corresponding contralateral muscle fiber. These differences can clearly be seen (Fig.3B.) in the ‘f’ muscles in females of any age, and ‘d’ and ‘e’ muscles in 4 week old males. This volume mapping will be useful in representing relative muscle volumes upon genetic manipulation or physical trauma.

### Specific Dorsal Longitudinal Muscles increase in volume differently over time in males and females

Female Drosophila are on an average are larger than male Drosophila. From our volume measurements, we plotted the total DLM volumes of females and males at different ages. In Fig.4A., total DLM volumes of females and males at days 2,7,14 and 28 days post eclosion have been plotted. Clearly, the average total DLM volumes of females is ∼50% larger than the male average total DLM volume, at all four time points. In both groups, an upward trend in total DLM volumes can clearly be seen over time. Indeed, total DLM volumes at day 28 are significantly larger than total DLM volumes at day 2 post eclosion (p<0.001) in both sexes.

We investigated how the volumes of muscles ‘a’, ‘b’, ‘c’, ‘d’, ‘e’ and ‘f’ change over time in males and females. Absolute volumes observed from individual muscles are plotted from the two groups in Fig.4B.

Each animal contributes two volumes of the same muscle from either side of the midline i.e. a^L^ and a^R^ are two independent observations from the same animal at the same time point. Thus, each left and right muscle count as separate volume observations from the same animal.

In Fig.4B., volumes of muscles ‘a’ to ‘f’ at day 2 post eclosion have been compared with volumes of muscles ‘a’ to ‘f’ at day 28 post eclosion respectively. Volumes of the muscles ‘b’, ‘c’ and ‘d’ from females show statistically significant increases over time, whereas in males, all muscles but ‘f’, show an increase in volumes in this time window. The reason for this sexually dimorphic difference in volumes, in very specific muscles is unclear.

### Individual DLM volumes are asymmetric on either side of the midline

The arrangement of muscles about the midline is symmetrical. If however, one muscle on one side of the midline, is to serve as an internal control for its corresponding contralateral muscle, a near complete symmetry of shape and volume may be expected. In order to quantify variation between volumes of the same muscle on either side of the midline, we calculated the ratio of left muscle volume to right muscle volume for every muscle fiber (‘a’ to ‘f’) in our entire data set. A left to right volume ratio of 1 would imply perfect volume symmetry in a muscle about the midline.

Fig.4C. shows box plots of individual left-to-right volume ratios for fibers ‘a’ to ‘f’, for females and males, at different times in adulthood. The average left to right muscle volume ratios, for all DLMs, in males and females tend towards unity. However, the spread of individual observations in these sets, ranges from below 0.75 to above 1.25.

Therefore, this variation in contralateral muscle volumes should be recognized while using them as internal controls.

No significant changes in left to right volume ratios was seen over time in any muscle group. Muscle volume often correlates with muscle deployment. This variation in left to right DLM volume ratios leads to an interesting idea, that there may exist a preference for the usage of one side over the other, in Drosophila. Evidence for bias in route choice and individual handedness has been presented in Honey bees(10).

### Non-myofibril-bundle volumes in Muscle fibers increase over time

It is possible, the separation of myofibril fascicles in wildtype DLMs as seen in Fig.1B. and Video 2 is an artefact of our MicroCT sample preparation. In such a case, it is to be expected that no consistent trend in the volume separating the fascicles may be seen over time. To test this hypothesis, we plotted the percentage of volume occupied by signal voxels (myofibril signal), in the three-dimensional region bounding each muscle.

In Fig.5. we plot the percent volume that myofibril signal occupies in each DLM over time in males and females. A consistent downward trend in these percent volume fractions occupied by myofibril signal in a muscle fiber can be seen. The volume fractions occupied by myofibril signal in muscles is reduced significantly between day 2 and day 28 post eclosion in all muscles in both males and females. The drop in average percent myofibril volume ranges from 5% to 15%. This consistent trend suggests this observation is unlikely to be an artefact.

The functional significance of this age dependent alteration in fascicle arrangement remains to be discovered. What anatomical, cellular and molecular features keep fascicles tightly bound initially but separate over time and how these contribute to flight, may inform our view on aging in muscles.

### Method applies to other insect species

Our interests lie largely in understanding adult Drosophila Dorsal Longitudinal Muscle development and homeostasis. However, the application of this method to other insect species may make an impact in comparative studies.

We tested our contrasting and scanning protocols on honeybee thoraces. Video 3 shows a MicroCT scan of an Apis species thorax from the vicinity of National Center for Biological Sciences. Dorso-Ventral Muscles and Dorsal Longitudinal muscle groups in the thorax can clearly be seen. The DLM group has been segmented out and clearly shows a vast difference in fiber number and shape from Drosophila DLMs.

The protocol was varied slightly for this prep: fixed thoraces were incubated in I_2_/KI solution for 72 hours, instead of 16 hours, to allow proper perfusion into the larger honeybee thorax. With appropriate variations in incubation times and I_2_/KI volumes, this protocol maybe applied to many other species.

We hope that our fixation and staining protocol and variations thereof will assist accurate *in situ* reporting of internal soft tissue across insect species.

## Materials and Methods

### Fly stocks

All volume measurements were performed on the *Canton S* strain of Drosophila.;; *UAS mito-GFP, UAS-mCD8RFP, mef2-Gal4* and *UAS-KDEL-RFP, UAS-mCD8GFP*; *mef2-Gal4* stocks were recombined from elements available in Bloomington Stock Center.

### Drosophila Husbandry

Wildtype *Canton S* flies were grown on corneal agar at 25 degrees centigrade on a 12 hour light dark cycle. For accurate and representative samples of aging, cohort of animals were collected within an hour post eclosion.

20 animals per vial was grown in previously mentioned conditions. Males and females were grown in separate vials.

### Immunohistochemistry

Adult thoraces were dissected in in 4% paraformaldehyde, 0.3% Triton-X diluted in phosphate buffered saline (PBS pH-7.5) and fixed for 10 mins per group, with the head, abdomen and legs removed. Each thorax was spread on a double sided taped glass slide, ventral side up. With a sharp blade, a section was made in the coronal plane. Samples were then subjected to three washes of 0.3% PTX (PBS + 0.3% Triton-X) for 15 mins each wash. Primary antibody staining was performed for overnight on a shaker at 4 degrees Celsius. After two 15 minute washes in PTX, samples were incubated in secondary antibodies with shaking for 4 hours at room temperature. After three 15 minute washes in in PTX, samples were mounted in Vectashield mounting media. For immunostaining, anti-GFP (Chick, 1:500, Abcam, Cambridge, UK), anti-RFP (Rabbit, 1:500, Abcam), Hoechst 33342 (1:500, ThermoFisher), Phalloidin (Alexa-647 conjugate, 1:500, ThermoFisher), were used. Secondary antibodies (1:500) from Invitrogen conjugated with Alexa fluor-488, 568 and 647 were used in immunostaining procedures.

### Sample preparation for MicroCT scanning

At appropriate time points, groups of flies were anesthetized on ice and transferred to 4% paraformaldehyde (PFA) made in PBS in a dish. Thoraces were dissected out by pulling outing heads and cutting away abdomens while retaining the wings. Thoraces were then transferred into a tube with 300 microliters of fixative solution, making sure all samples are submerged. These samples were incubated at room temperature for three hours with gentle shaking. Subsequently, fixative was aspirated out and discarded followed by two 15 min washes at room temperature with 1ml PBS per tube. 200 microliters of staining solution (1% elemental iodine with 2% potassium iodide dissolved in PBS) was then added to each tube making sure all thoraces were submerged. Samples were incubated in staining solution with gentle shaking at room temperature overnight.

### MicroCT scanning

Prior to scanning, each thorax washed twice in 500 microliters of PBS for 15 minutes at room temperature. Each thorax was dipped in paraffin oil to retain moisture during the scan. Individual thoraces were then mounted on a micro positioning stage tipped with petroleum jelly, and wings were used to position and stabilize the thorax. These samples were then scanned on Bruker Skyscan-1272 at 40kV, 250microA, 4940×3280 pixels without filters at 0.5micron resolution with a rotation step of .55 degrees.

## Data processing and volume calculation

From each sample, all projection images were imported to the NRecon software (Bruker Instruments) for a 3D reconstruction with 5 unit smoothing.

The 3D resolution of each CT virtual section is undersampled by a factor of 4, to speed up computation. The stack was reoriented in Dataviewer (Bruker) to align with coronal and sagittal planes. Regions of interest (ROI) specifying muscles were drawn on this reoriented stack in CTan (Bruker). Thresholding on the stack was done using the Ridler-Calvard method, and volumes of signal bearing voxels within the marked ROI were calculated from these binarized images.

Figs.4 and 5 were plotted using Python. Volume maps in Fig.3B. were manually drawn on Adobe Illustrator. Briefly, square roots of individual muscle volumes from Table 1 were calculated. All measurements were normalized to the smallest measurement in the entire dataset including both male and female measurements. Squares of corresponding side lengths in cm were drawn in Adobe Illustrator. The borders of each square were used to indicate spread in that measurement. The standard errors of mean were calculated from Standard deviations from the Table 1. Their square roots, normalized to the smallest measurement above, were used to determine border thickness in cm.

**Fig 4.**
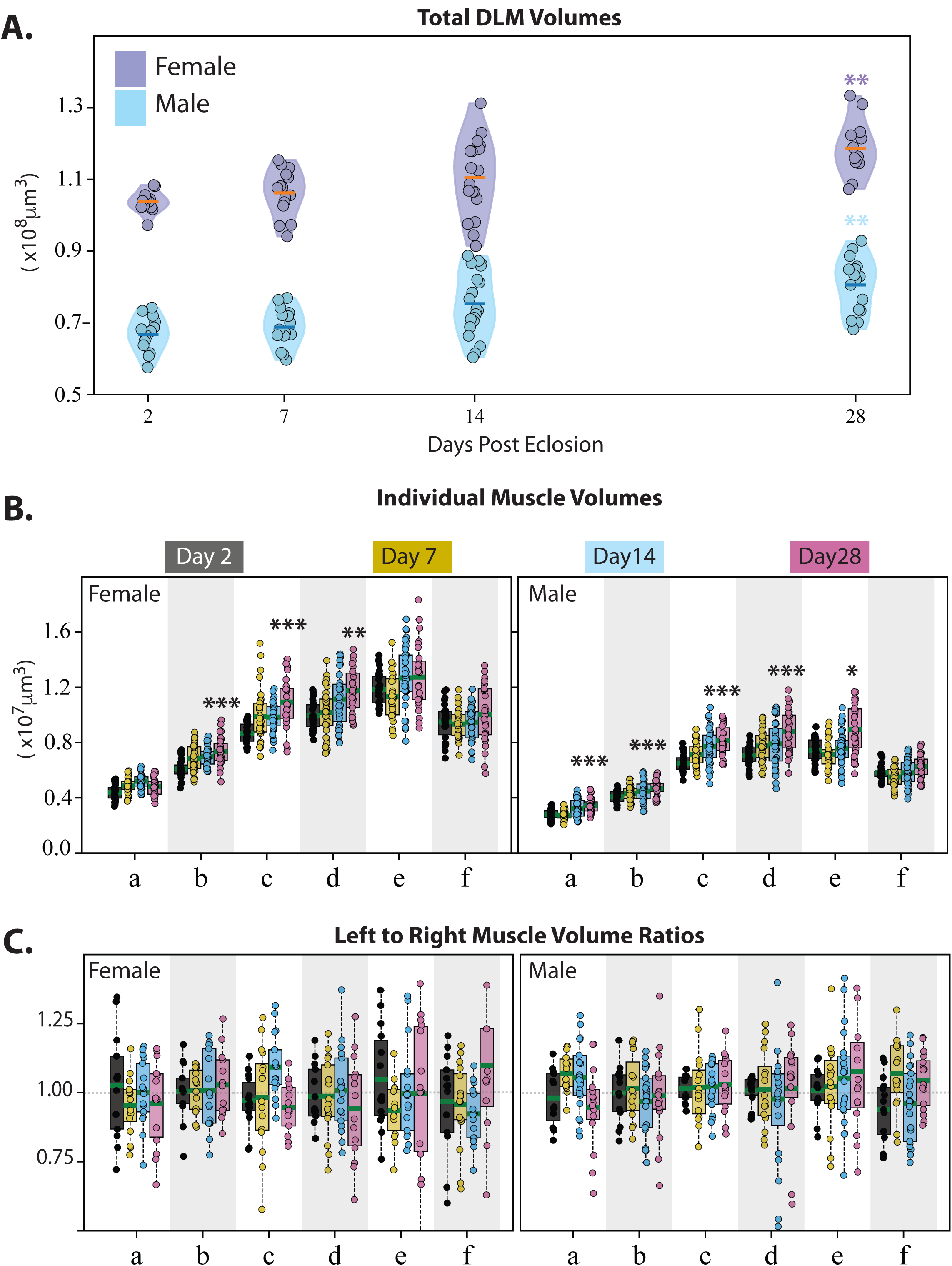
Variation in total and individual muscle volumes, and volume symmetry about the midline, for males and females over time. (A) Violin plots for Total DLM volumes of females (purple) and males (Blue) at days 2,7,14 and 28 post eclosion. Each point denotes an individual observation. Orange bars denote mean total DLM volumes in females and the blue bar denotes mean total DLM volumes in males. n=14-21 animals per sex per time point. ** denotes p<0.001 while comparing data from day 28 with day 2 using the Wilcoxon rank sum test. (B) Box plots for individual muscle volumes for ‘a’ to ‘f’ in females(left) and males (right) at days 2 (grey), 7 (Yellow),14 (blue) and 28 (Magenta) p.e. Each point denotes an individual observation. Green bars denote mean volumes per set. n=14-21 animals per sex per time point. *, **, *** denote p<0.01, <0.001, <0.0001 respectively while comparing data from day 28 with day 2 using the Wilcoxon rank sum test. (c) Box plots for Left muscle volume to Right muscle volume ratios, for muscles ‘a’ to ‘f’ in females(left) and males (right), at days 2 (grey), 7 (Yellow),14 (blue) and 28 (Magenta) p.e. Each point denotes an individual ratio. Green bars denote mean volumes per set. n=14-21 animals per sex per time point.

**Fig 5.**
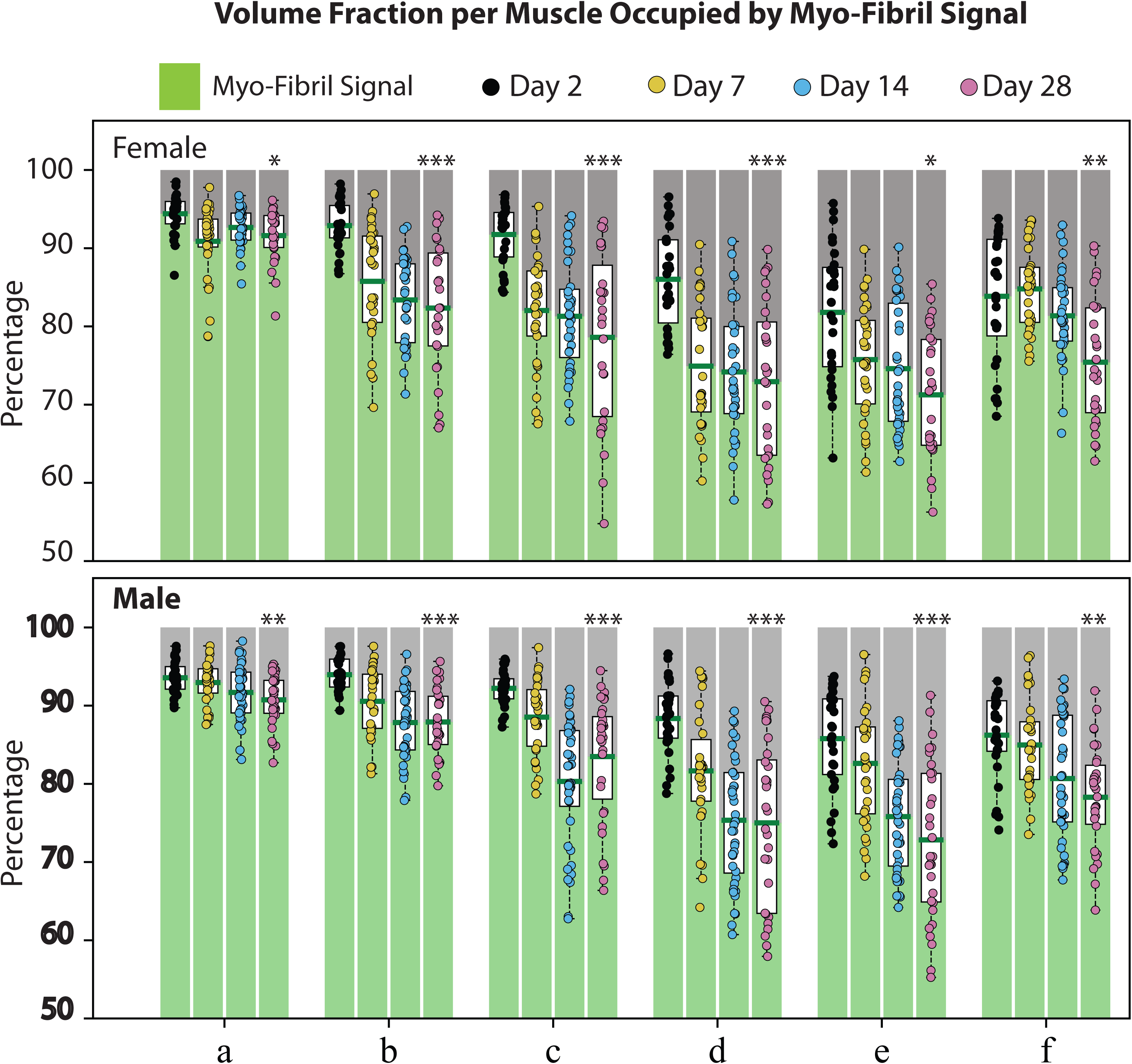
Variation in volume fraction occupied by myofibril signal for males and females over time. Box plots describing the percent of total volume of each DLM fibre, measured as in Fig.4a., that is occupied by non-zero signal (myo-fibril signal, green) and zero signal (grey). Measurements for females (top panel) and males (bottom panel), for muscles ‘a’ to ‘f’, at days 2 (black), 7 (Yellow),14 (blue) and 28 (Magenta) p.e. Each point denotes an individual observation. Green bars denote mean myo-fibril occupied volume percent, per set. n=14-21 animals per sex per time point. *, **, *** denote p<0.01, <0.001, <0.0001 respectively while comparing data from day 28 with day 2 using the Wilcoxon rank sum test.

## Discussion

In addition to assessing function, variations in tissue morphology through the processes of development, aging, disease conditions and repair, are critical to a full understanding organism function. Muscle function is key to quality of life, survival and metabolic regulation in many species(11-13). Drosophila muscles, especially DLMs, model homeostatic adult muscles with a fibrilar arrangement that is shared with mammalian skeletal muscles. The vast genetic toolkit, short lifespan and shared mechanisms of adult repair (14, 15), make Drosophila DLMs a promising model for adult human muscle repair and pathologies.

By employing a contrasting regime in near isotonic aqueous based medium in MicroCT scanning, we are able to measure *in situ* Drosophila muscle morphology and arrangement, at a significantly improved combination of scale and resolution. This protocol differs in the use of alcohol and high salt concentrations in buffers, that shrink soft tissue.

Through this protocol we have elucidated the variations in adult fruit fly DLM morphology in an aging and sex dependant manner.

We find that the Drosophila thorax is more fully packed with muscles as the animals age. While the total DLM volume increases with age in both males and females, not all DLM fibres grow in volume, and the fibres that do so, differ among males and females. At all ages, on average, volumes of left and corresponding right DLMs are symmetrical. However, individual measurements reveals a large variation from this mean. This is likely influenced by the packing of myofibrils within each muscle, which as we have shown, reduces in density with age in both sexes.

Many of our findings shine a light on aspects of DLM aging that couldn’t be satisfactorily interrogated with existing techniques. The use of MicroCT scanning with our protocol has made this possible. Further inquiries into the molecular bases of DLM growth, repair, myofibril fasciculation etc. can be pursued, based in these data. This is especially true of myopathy models where DLMs are likely to be distorted by the dissection process. We have also shown that this protocol is not limited to Drosophila and can be used in comparative studies across species. For instance, precise measurements of musculature in different castes of ant and honey bee colonies may further inform investigations in to the molecular details of their development (16).

In all, this technique promises to be a valuable addition to the toolkit of biology.

## Supporting information

Wildtype Thorax

Wildtype fascicles

Honey Bee Thorax

## Acknowledgements

This research was supported by NCBS and the Department of Science and Technology’s JC Bose Fellowship given to K. VijayRaghavan.

## Figure Legends

### Video1

#### Representative whole Drosophila thorax MicroCT scan and DLM segmentation

A whole Drosophila thorax with the head and abdomen dissected out. The thorax is positioned anterior to the left of the frame and posterior to the right at the start of the video. This orientation is reversed by second 5 of the video. At second 12, the thorax is oriented with the anterior side facing the viewer. At second 16, DLMs junctions on the cuticle are highlighted through red segmentation. DLMs (red) can be followed anterior to posterior in cross section up to second 30. The lateral planes of view are limited to mostly including DLMs by second 46. By the end of the video, the anterior posterior extension of DLMs on both sides of the midline can be seen.

### Video2

#### Continuous anterior to posterior extension of myo-fibril bundles (fascicles) in DLMs

The lateral planes of view limited to mostly including DLMs. At the start of the video, the thorax is oriented with the anterior to the right of the frame and posterior to the left. Separation of myofibril bundles within, and through the length of, each muscle can clearly be seen. By second 8, the thorax is oriented with the anterior side facing the viewer. The plane of view, then moves posterior-wards revealing continuous separation between myo-fibril bundles.

### Video3

#### Representative whole Honey bee MicroCT scan and DLM segmentation

A whole Honey bee thorax with the head and abdomen dissected out. The thorax is positioned anterior to the left and posterior to the right at the start of the video. At second 3, the thorax is oriented with the anterior side facing the viewer. The plane of view move anterior to posterior in cross sectional view up to second 13, with DLMs segment in red. Non DLM tissue has been removed from view between seconds 18 and 21.

